# Information Theory and Stem Cell Biology

**DOI:** 10.1101/116673

**Authors:** Rosanna C. G. Smith, Ben D. MacArthur

## Abstract

**Purpose of Review:** To outline how ideas from Information Theory may be used to analyze single cell data and better understand stem cell behaviour.

**Recent findings:** Recent technological breakthroughs in single cell profiling have made it possible to interrogate cell-to-cell variability in a multitude of contexts, including the role it plays in stem cell dynamics. Here we review how measures from information theory are being used to extract biological meaning from the complex, high-dimensional and noisy datasets that arise from single cell profiling experiments. We also discuss how concepts linking information theory and statistical mechanics are being used to provide insight into cellular identity, variability and dynamics.

**Summary:** We provide a brief introduction to some basic notions from information theory and how they may be used to understand stem cell identities at the single cell level. We also discuss how work in this area might develop in the near future.

## Introduction

Stem cells are characterized by their ability to self-renew and differentiate along multiple distinct lineages. Due to these remarkable properties there is much hope for stem cell based therapies in regenerative medicine. However, the development of such therapies will require a thorough understanding of the molecular mechanisms by which stem cells balance self-renewal and differentiation. Since stem cells are often rare (as in the adult) or exist only transiently (as in development), recent years have seen a growing focus on using single cell profiling technologies to understand stem cell dynamics. These studies have indicated that apparently functionally homogeneous stem cell populations can vary widely in their expression of important regulators of self-renewal and multipotency. In some cases this variability is driven by dynamic fluctuations of important master transcription factors, suggesting that stem cell heterogeneity has an important functional role [1–3]. However, the relationship between molecular heterogeneity and stem cell function are still not well understood.

Recent years have seen remarkable advances in single cell sequencing techniques, and it is now possible to profile large portions of the genome, or the entire transcriptome, in hundreds to thousands of individual cells in a single experiment [4–6]. Advances in single cell epigenetics and proteomics are not far behind [7–10]. These advances promise to transform our understanding of cellular identities, yet they also produce vast amounts of complex data, making it a significant challenge to distinguish meaningful biology from experimental noise. In the context of stem cell dynamics numerous reports have indicated that functionally homogeneous stem cell populations, both from the adult and the embryo, are highly heterogeneous with respect to their patterns of gene and protein expression [11–15]. However, the extent to which this variability plays a functional role, and the extent to which it represents variability due to inherent, but non-functional, expression noise are not clear. Therefore, in order to understand stem cell function at the individual cell level it has become increasingly necessary to use high-throughput profiling techniques to explore co-expression dynamics at the single cell level to identify rare (yet potentially functionally important) cells and determine how co-expression patterns change over time. The data provided by these experiments are fundamentally different from those obtained from measurements on cellular aggregates. While bulk methods typically provide estimates of the mean expression of each variable (e.g. gene) profiled over all cells in the aggregated sample (perhaps along with estimate of variance when the sample mean of multiple replicates are taken), they are not generally well suited to exploring dependencies between variables because they are only capable of examining expression patterns on average, not within individual cells. By contrast, since single cell methods profile co-expression patterns within individual cells they are able to provide a sample from the joint distribution of all the variables being profiled and so are much better suited to explore functional relationships between variables. Importantly, recent years have seen significant improvements in the efficiency of single cell RNA-seq methods, which now allow profiling of many tens of thousands of individual cells thereby improving estimates of joint expression distributions [6, 16, 17]. The experimental progress made in capturing multivariate single cell data has also stimulated research into new analysis techniques that are specifically designed to handle high-dimensional single cell data [18, 19]. These new analysis methods often make use of classical multivariate statistics and statistical approaches have provided insight into many stem cell systems including identification and characterization of mixtures of cellular states [20], comparison of different stem cell lines [21], rare cell identification [22] and cell lineage decision making [23]. However, methods from information theory are increasingly also being used to better understand how cellular expression patterns determine cellular identities.

## Information theory

Information theory has its roots in Shannon's work on communication and his famous 1948 paper laid out the mathematical theory of information [24, 25]. Shannon realized that in order to quantify the information content of a message it is necessary to consider the message's *context,* or how probable it is. An intuitive understanding of this can be seen in the following example. Consider a search for this article using only the last name of one of the authors. Which one is it best to choose? The knowledge that ‘Smith’ is a very common last name and ‘MacArthur’ is less common means that searching for ‘MacArthur’ is more likely to narrow the search and therefore likely to provide more information. The fact that ‘MacArthur’ is a more complex word than ‘Smith’ is irrelevant: it is the rarity of each name that dictates which to choose, not the name itself. In the context of gene expression, the fact that a cell has 7 transcripts of a particular mRNA does not in itself carry any information: this observation requires context in order to understand how much information is gained from the measurement. Without the context of how likely a read of 7 transcripts is, the information gained from the measurement is unknown (colloquially this is known as Shannon's zeroth law). So how do we calculate information gain? Shannon argued that any measure of information should satisfy three basic requirements: monotonicity, independence and branching. Monotonicity ensures that the information gained from a question with a wide variety of answers is greater than the information gained from the answer to a question with only a few possible answers. For example, to identify a specific person an answer to the question *“where do they come from*?” provides more information than an answer to the question *“are they female* ?”. Independence ensures that the total information gained from two independent questions should is a sum of the information gained from the questions separately. So, for example, the order in which the questions are asked should not matter. Lastly, branching ensures that when a series of questions is composed in a tree-like structure, the overall information gained by passing along a path through the tree is a weighted sum of the information gained from each branch point [25].

Shannon proved that the following function, which he called the entropy by analogy to the closely related thermodynamic entropy, uniquely satisfies these conditions. The Shannon entropy *H* is the expected amount of information gained from answering a question for which the probability of answer *x* is given by *p*(*x*),

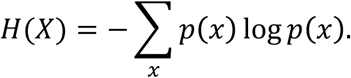

The entropy is a property of the probability distribution *p*(*x*), in the same way that the mean and variance are properties of *p*(*x*). Informally, the entropy is simply a measure of how “flat” or close to uniform *p*(*x*) is, and the “flatter” a distribution is, the greater the entropy and information gained. The units of entropy depend on the base of the logarithm: when the logarithm is taken to the base 2, as is common in information theory, entropy is measured in bits (one bit is the amount of information provided when observing one of two equally likely outcomes, e.g. the flip of a fair coin). Alternatively, entropy is measured in nats when using the natural logarithm (as is typically the case in statistical mechanics), and in hartleys when using base 10 (one hartley is the amount of information provided when observing one of 10 equally likely outcomes, e.g. a uniformly randomly chosen decimal digit). The equation for the entropy given above assumes that the random variable *X* is discrete. In practice many measures of interest, such as molecular concentrations, are continuous and the continuous analogue to the entropy above is known as the differential entropy [26]. In the discrete case, the entropy has some useful properties (for example, *H*(*X*) ≥ 0) that are not inherited by the differential entropy. To account for these differences, several closely related variations such as the Kullback-Leibler divergence (also known as the relative entropy) and its generalizations are commonly used to assess similarity between continuous expression distributions [27, 28]. For example, the widely-used t-SNE dimensionality reduction algorithm [29] (which has been used in several recent stem cell studies to explore heterogeneity in stem cell identities and cluster cell states [16, 17, 22, 23]) uses the Kullback-Leibler divergence to assess the similarity between the observed co-expression distribution and that obtained by projecting the data to a lower-dimensional space.

## Information theory and stem cell biology

The utility of the entropy in understanding cell identities many be illustrated by returning to our example of the measurement of 7 mRNA transcripts in a cell. To gain context to this reading, we need to better understand the natural variability of mRNA expression in the cell population of interest to determine how unusual this reading is. Consider the following two hypothetical scenarios for mRNA expression in a population of stem cells, as shown in Fig. 1A: (1) all cells in the population have 7 mRNA transcripts (i.e. 7 is the only answer to the question *how many transcripts are in the cell?* and occurs with probability 1). In this case, since all cells are the same with respect to their transcript counts, the observation of 7 transcripts cannot be used to discriminate one cell from another, and therefore does not impart any information. Accordingly, the entropy is *H =* −1 log(1) = 0 bits. (2) TWO stem cell subtypes are present in the population (types A and B). Cells of type A occur with probability 0 < *p* < 1 and have 7 transcripts, while cells of type B occur with probability (1 − *p*) and have zero transcripts. In this case, the observation of 7 transcripts allows us to positively discriminate cells of types A from those of type B and so imparts useful information. Furthermore, the amount of information we gain is related to the relative rarity of types A and B. In particular, the entropy is given by, *H =* −*p* log(*p*) − (1 − *p*) log (1 − *p*). Thus, when *p* is small, the observation of 7 transcripts in a cell is a rare event, but the observation of zero transcripts is a common event and so the entropy is low. Conversely, when *p* is large the observation of 7 transcripts is a common event while the observation of zero transcripts is a rare event and again the entropy is low. However, when cells of both types are common in the population (i.e if *p*~0.5) then the entropy reaches its maximum. In this example it is worth noting that the fact that cells of type B express no transcripts is not relevant to the calculation of entropy, they could have expressed any number of transcripts not equal to 7: all that is important is that cells of type B can be distinguished from those of type A by their mRNA transcript count.

**Figure 1:**
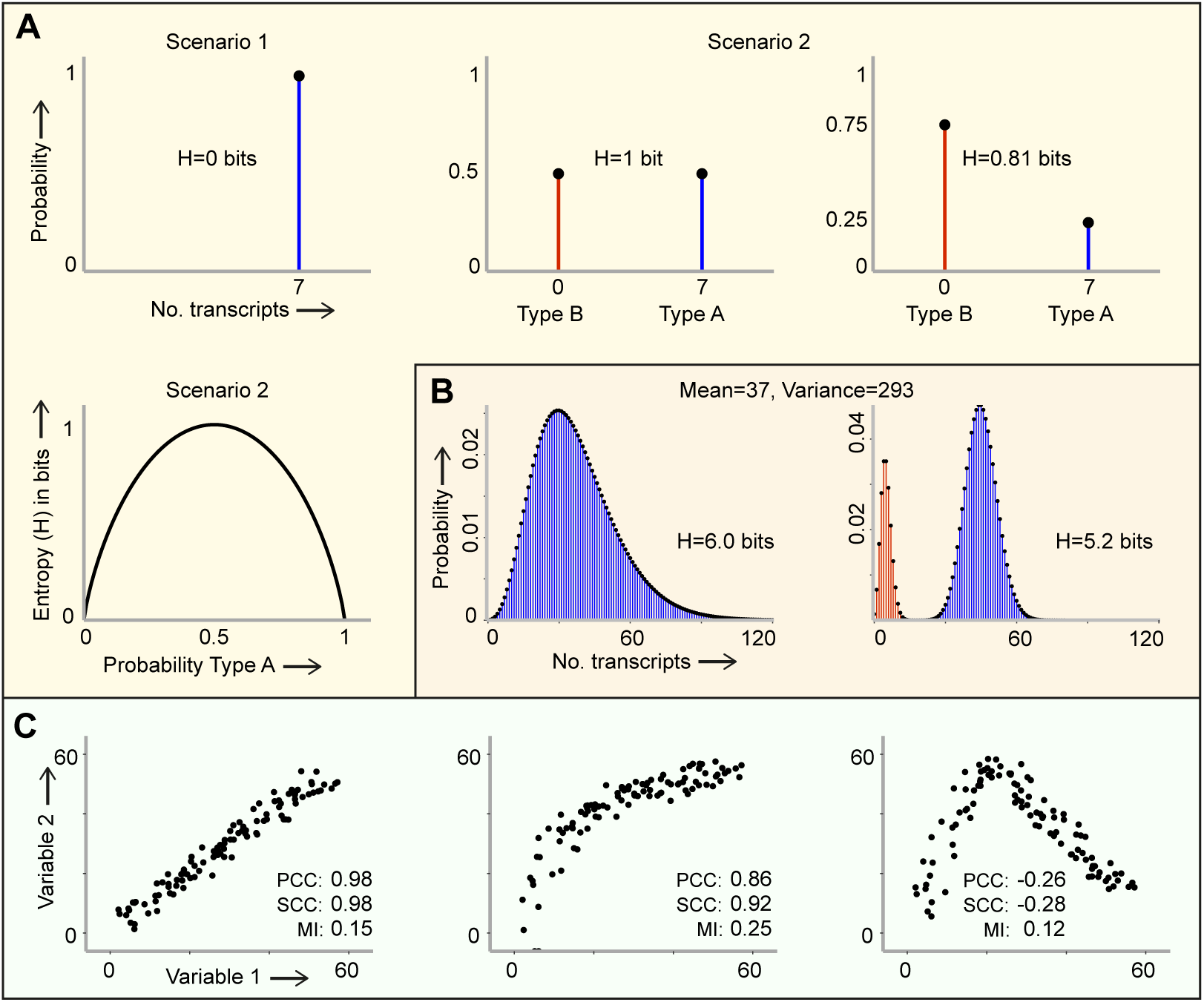
Entropy and Mutual Information. **A) Entropy of hypothetical binary cell types:** Scenario 1: All cells have 7 mRNA transcripts and entropy is zero (there is no uncertainty). Scenario 2: Cells are either type A (7 transcripts), which occurs with probability *p*, or type B (zero transcripts), which occurs with probability 1 − *p*. When there is an equal probability of observing either cell type (*p* = 0.5), we are maximally uncertain about the identity of a randomly drawn cell and the entropy *H =* 1 bit, the same as for tossing a fair coin. When there are unequal probabilities, for example when *p* = 0.25, uncertainty is reduced the entropy is less than 1 bit. The final panel gives the relationship between entropy and *p* from which it can be seen that maximum entropy occurs when *p* = 0.5. **B) Entropy of distributions:** Distributions of transcript abundance are typically not binary, but rather exhibit a spread of possible outcomes. Examples of a unimodal and a bimodal distribution with the same mean and variance, but different entropies are shown. In the unimodal case the measures such as the mean and variance may make good sense. However, in the bimodal case the population mean is not characteristic of either of the two subpopulations (it is rare to find a cell with the mean level of expression) and the variance as a measure of the spread about this mean is also misleading. By contrast the entropy, which measures the amount of uncertainty we have concerning the identity of a randomly draw cell from the population, provides useful information about cell-cell variability. **C) Mutual information as a measure of association:** Association between two random variables can be assessed by Pearson's correlation coefficient (PCC), which considers the strength of linear association, Spearman's correlation coefficient (SCC) which is based on rankings, and mutual information (MI) which assesses how much information one variable provides about the other. All three measures can assess linear associations well (left panel), SCC is a good measure of non-linear, monotonic associations (middle panel), but neither PCC or SCC are good measures of association for non-linear, non-monotonic associations (right panel). However, the MI may be used to determine that the two variables are related.

In practice we would not expect that all cells express a given mRNA at one of two fixed levels: rather, intrinsic noise in gene expression naturally gives rise to variations in gene expression levels over time within each individual cell, and within the cell population at any fixed time (see Fig. 1B). While it cannot often be calculated explicitly as above, the entropy can nevertheless be estimated from experimental data to better understand this natural variation, (it should be noted that entropy estimation is subject some technical issues including the effect of data binning and bias on entropy estimation [30–32]). For example, it has been suggested that a high degree of cell-cell variability in gene expression patterns within a functionally pure population, as quantified by the entropy of the joint expression distribution, is characteristic of undifferentiated pluripotent cells [33–35]. Similarly, by considering patterns of gene expression in light of a known signaling networks, Teschendorff and colleagues have argued that both pluripotent cells and cancer cells are associated with a state of high network entropy, characterized by the promiscuous co-expression of important hub proteins [36–38]. Relatedly, it has been observed that the entropy of gene expression developing tissues increases with time in a manner that is closely related to differentiation dynamics [39–41].

While the entropy is good at assessing how likely it is that a particular expression value will occur, it is not well suited to assessing relationships between co-expression patterns. TO do so a related measure, the mutual information (MI), is also widely used. Consider 2 discrete random variables, *X* and *Y,* which may be related in some unknown way. The entropy of the joint probability density *p*(*x*, *y*) is:

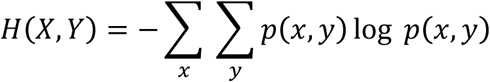

Informally this is a measure of the information content of the joint distribution, but it is not a direct measure of association between the two random variables. In order to assess whether one variable provides information about the other, the mutual information *I* (*X; Y*) may be used [26]. The mutual information compares the observed joint probability density with that which would be observed if the two random variables were independent. In particular

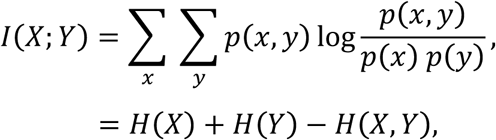

where *H*(*X*) and *H*(*Y*) are the marginal entropies. If *X* and *Y* are independent then *p*(*x*,*y*) *= p*(*x*) *p*(*y*), so 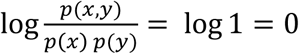 for all *x* and *y* and therefore *I*(*X; Y*) *=* 0. In this case, knowledge of one variable does not provide any information about the other variable. More generally since *I*(*X*; *Y*) *=* I(*Y; X*) ≥ 0 the magnitude of the MI is a measure of the extent to which the observed joint distribution deviates from independence: larger values of MI indicate a stronger dependency between *X* and *Y.* The advantage of MI as a measure of association is that does not specify in advance the nature of the relationship between *X* and *Y* so it can capture non-linear, non-monotonic, dependencies between variables in a general way that traditional correlation measures cannot (see Fig. 1C for some examples).

Since the mutual information assesses the extent to which two random variables are independent of one another, it can be used to identify putative functional relationships between experimentally observed variables (e.g. genes or proteins) [42, 43]. For this reason, there has been much interest in using information-theoretic methods to infer genetic regulatory networks from gene expression data, in order to better understand cellular dynamics. Typically these methods make use of generalizations of the MI as well as more advanced information theory concepts to assess co-dependencies between multiple variables [44–46]. Recent examples include the use of: the data processing inequality [47, 48]; the conditional mutual information, which assess the dependency between 2 random variables, conditioned on a third [49]; information-theoretic redundancy, which assesses the extent to which observed distributions deviate from the maximum entropy uniform distribution [50] and combinations of these measures [51, 52]. In the context of stem cell biology information-based network reconstruction methods have been used with some success to identify novel regulators of pluripotency and lineage specifiers [53, 54], as well as track changes in network structures during cellular differentiation [52, 55].

## Conclusions

Here we have summarized some of the ways that information theory can be used in combination with multivariate statistics to investigate stem cell identities. Although information-theoretic measures are not always intuitive and their practical application needs careful consideration, information theory provides a suite of tools that can help make the most of experimentally hard-earned data. As well as providing improved measures of variability and association, information theory also has a natural relationship with statistical mechanics [56, 57], and thereby provides a natural approach to the investigation of cellular dynamics. Statistical mechanics addresses the question of how observable ‘macroscopic’ properties of a system arise from unobserved ‘microscopic’ dynamics. For example, the pressure of a gas in a confined container (a macrostate) depends upon the average kinetic energy of the molecules in the gas and can therefore be predicted without detailed knowledge of the instantaneous position and velocity of all the individual gas molecules involved (a microstate). In the 1950s Jaynes showed that statistical mechanics could be derived directly from information-theoretic principles [56, 57]. For example, he observed that the Boltzmann distribution, which is ubiquitous in statistical mechanics, arises naturally as the maximum entropy probability distribution subject to appropriate physical constraint. It would be interesting to see if similar approaches can be used to better understand cell-cell variability in stem cell systems: do observed patterns of variability in stem cell populations reflect natural biological constraints? If so, what are they? To what extent does cell-cell variability relate to stem cell function? Can a general theory of regulated cellular variability be derived using physical and information-theoretic principles? Some minor progress has been made towards these aims [33, 58–60] and this is an exciting area of current research, yet there is still much to be done. Although the relationships between cell-cell variability, entropy and cell function have yet to be fully deciphered, ongoing research indicates that information-theoretic measures can provide insight into cellular identities that are not apparent from more traditional multivariate statistical methods. We anticipate that advances in the accuracy and reductions in the cost of single cell methods are likely to see increased interest in the development and use of these methods in the near future.

## Acknowledgements

This work was funded by BBSRC Grant No. BB/L000512/1.

## References

1. Semrau S, van Oudenaarden A (2015) Studying lineage decision-making in vitro: emerging concepts and novel tools. Annu Rev Cell Dev Biol 31:317–45

2. Moignard V, Göttgens B (2016) Dissecting stem cell differentiation using single cell expression profiling. Curr Opin Cell Biol 43:78–86

3.** Kumar P, Tan Y, Cahan P (2017) Understanding development and stem cells using single cell-based analyses of gene expression. Development 144:17–32 Review of single cell transcriptome analysis methods and in their application in stem cell and developmental biology.

4.** Grün D, Van Oudenaarden A (2015) Design and Analysis of Single-Cell Sequencing Experiments. Cell 163:799–810 A thorough review of the recent advances in single cell transcriptome sequencing; comparison of preparation and sequencing methods and analysis techniques.

5. Kolodziejczyk AA, Kim JK, Svensson V, Marioni JC, Teichmann SA (2015) The Technology and Biology of Single-Cell RNA Sequencing. Mol Cell 58:610–620

6. Ziegenhain C, Vieth B, Parekh S, Reinius B, Guillaumet-Adkins A, Smets M, Leonhardt H, Heyn H, Hellmann I, Enard W (2017) Comparative Analysis of Single-Cell RNA Sequencing Methods: Molecular Cell. 631–643

7. Bendall SC, Simonds EF, Qiu P, et al (2011) Single-Cell Mass Cytometry of Differential Immune and Drug Responses Across a Human Hematopoietic Continuum. Science (80-) 332:687–696

8. Cusanovich DA, Daza R, Adey A, Pliner HA, Christiansen L, Gunderson KL, Steemers FJ, Trapnell C, Shendure J (2015) Epigenetics. Multiplex single-cell profiling of chromatin accessibility by combinatorial cellular indexing. Science 348:910–4

9. Spitzer MH, Nolan GP (2016) Mass Cytometry: Single Cells, Many Features. Cell 165:780–791

10. Budnik B, Levy E, Slavov N (2017) Mass-spectrometry of single mammalian cells quantifies proteome heterogeneity during cell differentiation. bioRxiv 102681

11. Hatano S-Y, Tada M, Kimura H, Yamaguchi S, Kono T, Nakano T, Suemori H, Nakatsuji N, Tada T (2005) Pluripotential competence of cells associated with Nanog activity. Mech Dev 122:67–79

12. Chambers I, Silva J, Colby D, Nichols J, Nijmeijer B, Robertson M, Vrana J, Jones K, Grotewold L, Smith A (2007) Nanog safeguards pluripotency and mediates germline development. Nature 450:1230–1234

13. Hayashi K, Lopes SMC de S, Tang F, Surani MA (2008) Dynamic equilibrium and heterogeneity of mouse pluripotent stem cells with distinct functional and epigenetic states. Cell Stem Cell 3:391–401

14. Toyooka Y, Shimosato D, Murakami K, Takahashi K, Niwa H (2008) Identification and characterization of subpopulations in undifferentiated ES cell culture. Development 135:909–918

15. Canham MA, Sharov AA, Ko MSH, Brickman JM (2010) Functional heterogeneity of embryonic stem cells revealed through translational amplification of an early endodermal transcript. PLoS Biol 8:e1000379

16.* Macosko EZ, Basu A, Satija R, et al (2015) Highly Parallel Genome-wide Expression Profiling of Individual Cells Using Nanoliter Droplets. Cell 161:1202–1214 Together with the study by Klein et. al, this study on droplet-based single cell RNA sequencing was a breakthrough method, making single cell transcriptome data more accessible.

17.* Klein AM, Mazutis L, Akartuna I, Tallapragada N, Veres A, Li V, Peshkin L, Weitz DA, Kirschner MW (2015) Droplet Barcoding for Single-Cell Transcriptomics Applied to Embryonic Stem Cells. Cell 161:1187–1201 Together with the study by Macosko et. al, this study on droplet-based single cell RNA sequencing was a breakthrough method, making single cell transcriptome data more accessible to researchers.

18. Kumar RM, Cahan P, Shalek AK, et al (2014) Deconstructing transcriptional heterogeneity in pluripotent stem cells. Nature 516:56–61

19. Bacher R, Kendziorski C (2016) Design and computational analysis of single-cell RNA-sequencing experiments. Genome Biol 17:63

20. Kolodziejczyk AA, Kim JK, Tsang JCH, et al (2015) Single Cell RNA-Sequencing of Pluripotent States Unlocks Modular Transcriptional Variation. Cell Stem Cell 17:471–485

21. Yan L, Yang M, Guo H, et al (2013) Single-cell RNA-Seq profiling of human preimplantation embryos and embryonic stem cells. Nat Struct Mol Biol 20:1131–9

22. Grün D, Lyubimova A, Kester L, Wiebrands K, Basak O, Sasaki N, Clevers H, van Oudenaarden A (2015) Single-cell messenger RNA sequencing reveals rare intestinal cell types. Nature 525:251–5

23. Olsson A, Venkatasubramanian M, Chaudhri VK, Aronow BJ, Salomonis N, Singh H, Grimes HL (2016) Single-cell analysis of mixed-lineage states leading to a binary cell fate choice. Nature 537:698–702

24. Shannon CE (1948) A mathematical theory of communication. Bell Syst Tech J 27:379–423

25. Bialek W (2012) Biophysics: Searching for Principles. Princeton University Press, Oxford, UK

26. Cover TM, Thomas JA (2005) Elements of Information Theory, Second. Elem Inf Theory. doi: 10.1002/047174882X

27. Kullback S, Leibler RA (1951) On Information and Sufficiency. Ann Math Stat 22:79–86

28. Tonge PD, Olariu V, Coca D, Kadirkamanathan V, Burrell KE, Billings SA, Andrews PW (2010) Prepatterning in the stem cell compartment. PLoS One 5:e10901

29. Van Der Maaten LJP, Hinton GE (2008) Visualizing high-dimensional data using t-sne. J Mach Learn Res 9:2579–2605

30. Olsen C, Meyer PE, Bontempi G (2009) On the impact of entropy estimation on transcriptional regulatory network inference based on mutual information. EURASIP J Bioinform Syst Biol 2009:308959

31. Hausser J, Strimmer K (2009) Entropy inference and the James-Stein estimator, with application to nonlinear gene association networks. J Mach Learn Res 10:1469–1484

32. McMahon SS, Lenive O, Filippi S, Stumpf MPH (2015) Information processing by simple molecular motifs and susceptibility to noise. J R Soc Interface 12:597

33. MacArthur BD, Lemischka IR (2013) Statistical mechanics of pluripotency. Cell 154:484–489

34.* Grün D, Muraro MJ, Boisset JC, et al (2016) De Novo Prediction of Stem Cell Identity using Single-Cell Transcriptome Data. Cell Stem Cell 19:266–277 This study incorporates the use of transcriptome entropy in identification of stem cells in mixed populations and is biomedically relevant in many systems where stem cells are not currently well characterised.

35. Guo M, Bao EL, Wagner M, Whitsett JA, Xu Y (2016) SLICE: Determining cell differentiation and lineage based on single cell entropy. Nucleic Acids Res. doi: 10.1093/nar/gkw1278

36. Teschendorff AE, Severini S (2010) Increased entropy of signal transduction in the cancer metastasis phenotype. BMC Syst Biol 4:104

37. West J, Bianconi G, Severini S, Teschendorff AE (2012) Differential network entropy reveals cancer system hallmarks. Sci Rep 2:802

38. Banerji CRS, Miranda-Saavedra D, Severini S, Widschwendter M, Enver T, Zhou JX, Teschendorff AE (2013) Cellular network entropy as the energy potential in Waddington's differentiation landscape. Sci Rep 3:3039

39. Anavy L, Levin M, Khair S, Nakanishi N, Fernandez-Valverde SL, Degnan BM, Yanai I (2014) BLIND ordering of large-scale transcriptomic developmental timecourses. Development 141:1161–1166

40. Piras V, Tomita M, Selvarajoo K (2014) Transcriptome-wide Variability in Single Embryonic Development Cells. Sci Rep 4:7137

41. Richard A, Boullu L, Herbach U, et al (2016) Single-cell-based analysis highlights a surge in cell-to-cell molecular variability preceding irreversible commitment in a differentiation process. PLOS Biol 14:e1002585

42. Antebi YE, Reich-Zeliger S, Hart Y, Mayo A, Eizenberg I, Rimer J, Putheti P, Pe’er D, Friedman N (2013) Mapping differentiation under mixed culture conditions reveals a tunable continuum of T cell fates. PLOS Biol 11:e1001616

43. Smith RCG, Stumpf PS, Ridden SJ, Sim A, Filippi S, Harrington H, MacArthur BD (2016) Nanog fluctuations in ES cells highlight the problem of measurement in cell biology. bioRxiv 60558

44. Allen JD, Xie Y, Chen M, Girard L, Xiao G (2012) Comparing statistical methods for constructing large scale gene networks. PLOS One 7:e29348

45. Villaverde AF, Ross J, Banga JR (2013) Reverse engineering cellular networks with information theoretic methods. Cells 2:306–29

46.** Mc Mahon SS, Sim A, Filippi S, Johnson R, Liepe J, Smith D, Stumpf MPH (2014) Information theory and signal transduction systems: From molecular information processing to network inference. Semin Cell Dev Biol 35:98–108 Review of information theory measures for inferring gene regulatory networks, including discussion of discretization and entropy estimator methods.

47. Nemenman I, Basso K, Wiggins C, Stolovitzky G, Favera RD, Califano A (2004) ARACNE: An Algorithm for the Reconstruction of Gene Regulatory Networks in a Mammalian Cellular Context. BMC Bioinformatics 7:1471–2105

48. Basso K, Margolin A, Stolovitzky G, Klein U, Riccardo D-F, Califano A (2005) Reverse engineering of regulatory networks in human B cells. Nat Genet 37:382–390

49. Liang K-C, Wang X (2008) Gene regulatory network reconstruction using conditional mutual information. EURASIP J Bioinform Syst Biol 2008:253894

50. Meyer PE, Kontos K, Lafitte F, Bontempi G (2007) Information-theoretic inference of large transcriptional regulatory networks. EURASIP J Bioinforma Syst Biol 2007:79879

51. Villaverde AF, Ross J, Morán F, Banga JR (2014) MIDER: Network inference with mutual information distance and entropy reduction. PLoS One 9:e96732

52. Chan TE, Stumpf MPH, Babtie AC (2016) Multivariate Information Measures. bioRxiv. doi: https://doi.org/10.1101/082099

53. Kushwaha R, Jagadish N, Kustagi M, et al (2015) Interrogation of a context-specific transcription factor network identifies novel regulators of pluripotency. Stem Cells 33:367–377

54. Okawa S, Angarica VE, Lemischka I, Moore K, del Sol A (2015) A differential network analysis approach for lineage specifier prediction in stem cell subpopulations. Syst Biol Appl 1:15012

55. Stumpf PS, Smith RCG, Lenz M, et al (2017) Stem cell differentiation is a stochastic process with memory. bioRxiv 101048

56. Jaynes ET (1957) Information theory and statistical mechanics. Phys Rev 106:620–630

57. Jaynes ET (1957) Information theory and statistical mechanics. II. Phys Rev 108:171–190

58. Garcia-Ojalvo J, Martinez Arias A (2012) Towards a statistical mechanics of cell fate decisions. Curr Opin Genet Dev 22:619–26

59. Ridden SJ, Chang HH, Zygalakis KC, MacArthur BD (2015) Entropy, Ergodicity, and Stem Cell Multipotency. Phys Rev Lett 115:208103

60. Moris N, Pina C, Arias AM (2016) Transition states and cell fate decisions in epigenetic landscapes. Nat Rev Genet 17:693–703

